# SOX9 promotes stress-responsive transcription of VGF nerve growth factor inducible gene in kidney epithelial cells

**DOI:** 10.1101/2020.07.06.189829

**Authors:** Ji Young Kim, Yuntao Bai, Laura A. Jayne, Ferdos Abdulkader, Megha Gandhi, Samir V. Parikh, Min-Ae Song, Amandeep Bajwa, Navjot Singh Pabla

## Abstract

Acute kidney injury (AKI) is a common clinical condition associated with diverse etiologies and abrupt loss of renal function. In patients with sepsis, rhabdomyolysis, cancer, as well as cardiovascular disorders, the underlying disease or associated therapeutic interventions can cause hypoxic, cytotoxic, and inflammatory insults to renal tubular epithelial cells (RTECs) resulting in the onset of AKI. To uncover stress-responsive disease-modifying genes, here we have carried out renal transcriptome profiling in three distinct murine models of AKI. We find that Vgf nerve growth factor inducible gene upregulation is a common transcriptional stress response in RTECs to ischemia, cisplatin, and rhabdomyolysis-associated renal injury. The Vgf gene encodes a secretory peptide precursor protein that has critical neuro-endocrine functions; however, its role in the kidneys remains unknown. Our functional studies show that RTEC-specific Vgf gene ablation exacerbates ischemia, cisplatin, and rhabdomyolysis-associated AKI in vivo and cisplatin-induced RTEC cell death in vitro. Importantly, addback experiments showed that aggravation of cisplatin-induced renal injury caused by Vgf gene ablation is partly reversed by TLQP-21, a Vgf-derived peptide. Finally, in vitro and in vivo mechanistic studies showed that injury-induced Vgf upregulation in RTECs is driven by the transcriptional regulator Sox9. These findings reveal a crucial downstream target of the Sox9-directed transcriptional program and identify Vgf as a stress-responsive protective gene in kidney epithelial cells.

## Introduction

Acute kidney injury (AKI) is a heterogeneous clinical syndrome that is associated with adverse short and long-term sequelae (1). AKI usually occurs in the setting of other diseases, such as sepsis (2), rhabdomyolysis (3), cardiovascular (4) and oncological diseases (5), where the underlying disease and or associated therapy cause abrupt loss of renal function. As a result, the pathophysiology of AKI is generally complex due to the existence of multiple etiologies such as the presence of sepsis, ischemia, and therapy-induced nephrotoxicity (6). AKI-associated mortality depends on the severity and can be significantly high in critically ill patients (7). Importantly, patients who survive an episode of AKI are at increased risk for major adverse cardiovascular events, as well as for progression to chronic kidney disease (8).

Disorders such as sepsis, cancer, rhabdomyolysis as well as therapeutic interventions such as cardiac surgery and chemotherapy are associated with inflammatory, toxic, and hypoxic insults to renal tubular epithelial cells (RTECs). The resulting RTEC dysfunction and cell death are the hallmarks of AKI (9). RTEC dysfunction and renal impairment clinically manifest as systemic electrolyte and fluid imbalances along with accumulation of metabolic waste, which can trigger multi-organ failure (7). The pathogenesis of AKI is multifaceted due to the involvement of various intracellular pathways (6,10–12) in RTEC dysfunction and cell death (9) as well as the contribution of vascular (13–15) and immune cells (16,17) in renal impairment.

Both the etiology and pathophysiology of AKI is complex. To identify common stress responsive genes, we have carried out genome-wide transcriptome analysis in three mouse models of AKI. Our studies identify the nerve growth factor-inducible gene, Vgf (nonacronymic; unrelated to VEGF) as a stress-responsive gene that is upregulated during ischemic, nephrotoxic, and rhabdomyolysis-associated kidney injury. Vgf was originally identified as a nerve growth factor (Ngf) - inducible gene in a neuroendocrine cell line and is expressed in specific neurons and endocrine cells in the brain and periphery (18). Vgf gene encodes a precursor polypeptide, which is proteolytically cleaved to generate several bioactive peptides, the best studied of which are TLQP-2, TLQP-62, and AQEE-30 (19). In the central nervous system, the secreted Vgf-derived peptides regulate neuronal activity, survival and progenitor proliferation (20–22). Furthermore, germline Vgf knockout mice have significantly reduced body weight, increased energy expenditure, and are resistant to diet-induced obesity, indicating that Vgf-derived peptides are critical regulators of energy homeostasis (20,23).

Interestingly, the role of Vgf in kidney physiology and pathophysiology remains unknown. Here, using transcriptome profiling and RTEC-specific gene ablation studies, we report that Vgf is a stress inducible gene that plays a protective role during the development of AKI.

## Results

### Mouse models of acute kidney injury

To identify common stress-induced cellular transcriptome changes linked to the pathogenesis of acute kidney injury, we sought to perform bulk RNA sequencing of renal tissues from distinct murine models of AKI. To this end, we utilized the well-characterized mouse models of ischemia-reperfusion injury (IRI), drug-induced nephrotoxicity (cisplatin), and rhabdomyolysis-mediated kidney injury. IRI-associated AKI results from a generalized or localized impairment of oxygen and nutrient delivery to the kidneys (6). Cisplatin nephrotoxicity results from specific drug uptake (24) and direct toxicity to tubular epithelial cells. Rhabdomyolysis-associated AKI results from skeletal muscle injury and the subsequent myoglobin release into the systemic circulation, which causes renal dysfunction (3).

In these mouse models, bilateral ischemic surgery, intra-peritoneal cisplatin injection, and intramuscular glycerol injection trigger AKI within 24-72 hours. The development and progression of AKI was determined by accumulation of nitrogenous waste (blood urea nitrogen and serum creatinine) and histological analysis of tissue damage (H&E staining and renal damage score). During ischemia (**Fig. 1A-C**) and rhabdomyolysis-associated (**Fig. 1D-F**) kidney injury, onset of renal impairment occurs 24 hours post-surgery or -injection, while in the cisplatin-associated kidney injury models, renal impairment is observed 72 hours post-injection (**Fig. 1G-I**). Histological analysis revealed similar tubular damage in IRI (24 hours), rhabdomyolysis (24 hours) and cisplatin (72 hours) groups (**Fig. 1J**).

**Figure 1:**
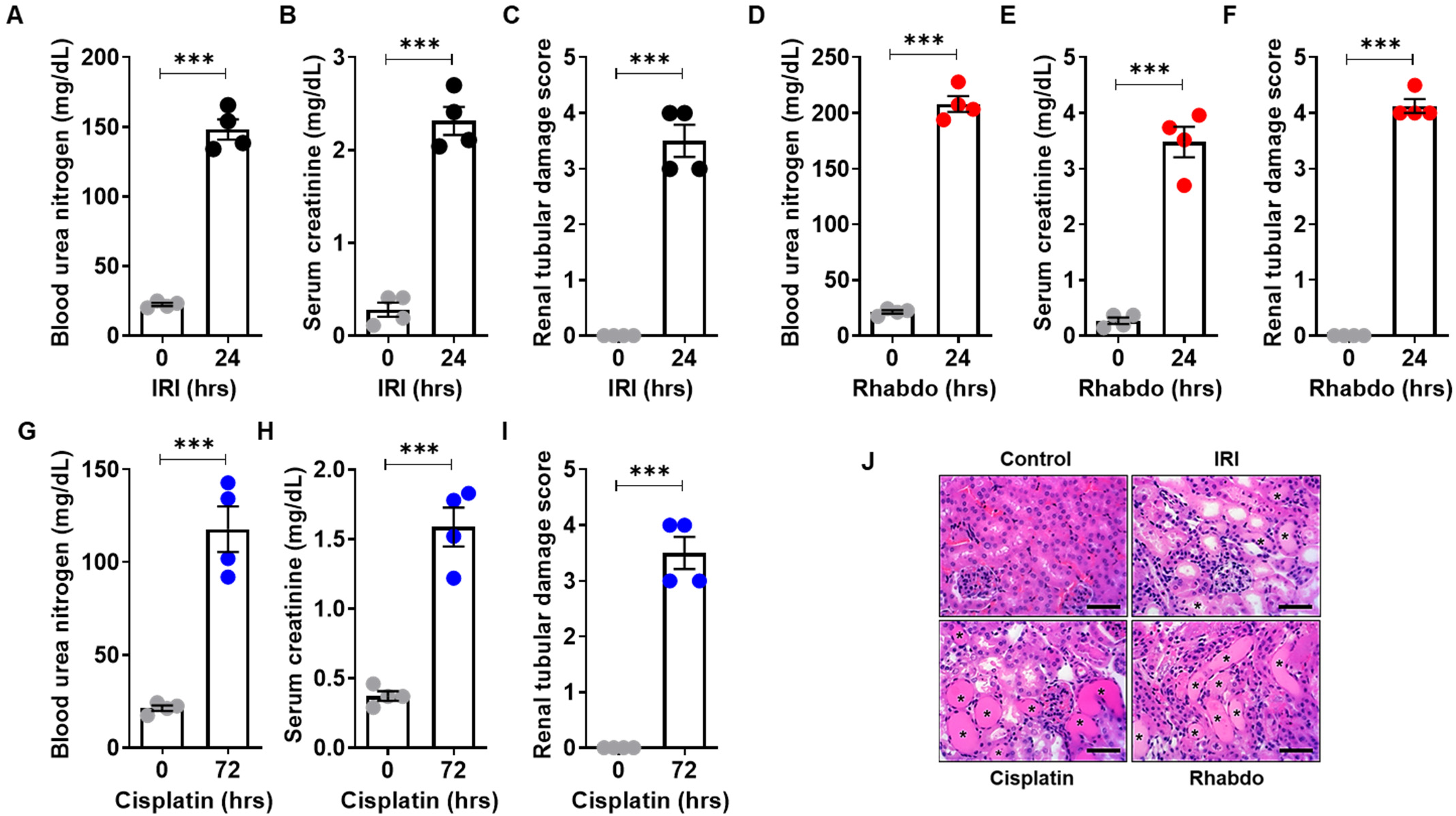
Mouse models of acute kidney injury. Ischemic, nephrotoxic, and rhabdomyolysis-associated acute kidney injury was induced in 8-12 weeks old male C57BL/6J mice. Bilateral renal ischemia (30 minutes) was induced for 30 min followed by reperfusion for 24 hours. Blood urea nitrogen (**A**), serum creatinine (**B**), and histological analysis (**C**) were performed to examine renal function and damage. Rhabdomyolysis was induced in male mice by glycerol injection (7.5 ml/kg 50% glycerol) in the hind-leg muscles followed by measurement of renal function (**D-E**) and histological analysis (**F**) of renal damage. Nephrotoxicity was induced in mice by a single intraperitoneal injection of cisplatin (30 mg/kg), followed by BUN (**G**), serum creatinine (**H**), and histological analysis (**I**) at the indicated time-points. (**J**) Representative H&E staining depicting renal tubular damage (indicated by an asterisk) linked with ischemia, cisplatin-nephrotoxicity, and rhabdomyolysis associated AKI. The graphs (A-I,n=4) are representative of two independent experiments. In all the bar graphs, experimental values are presented as mean ± s.e.m. The height of error bar = 1 s.e. and p < 0.05 was indicated as statistically significant. . Student’s t-test (A-I) was carried out and statistical significance is indicated by *p < 0.05, **p < 0.01, ***p < 0.001. Scale bar (J): 100 μm.

### Transcriptome profiling of AKI-associated differentially expressed genes

Due to temporal differences in the onset of kidney injury, we chose to compare gene expression signals at time-points where the extent of kidney injury is similar in the three groups. To this end, we isolated renal tissues from control (mock and vehicle, n=8), IRI (24 hours, n=4), rhabdomyolysis (24 hours, n=4), and cisplatin (72 hours, n=4) treated mice and then performed RNA sequencing (4-8 biological replicates). Principal component analysis (PCA) showed that the biological replicates clustered together across groups, signifying a high degree of similarity (**Fig. 2A**). Hierarchical clustering (**Fig. 2B**) revealed both divergent and convergent gene signatures between control and the three AKI groups. In the three AKI conditions (FDR<0.05 and fold change≥2), we identified a common set of 1501 differentially (771 genes were downregulated and 709 genes were upregulated) expressed genes (**Fig. 2C**). In **Supplementary Files 1** and **2**, we have provided the complete list and normalized expression levels of all detected and differentially expressed genes. Enrichment of genes related to glutathione, nicotinamide, and fatty acid metabolism was observed upon gene ontology (GO) and KEGG pathway analysis **(Suppl. File 3)**. These pathways have been recently probed for their role in renal dysfunction (25,26).

**Figure 2:**
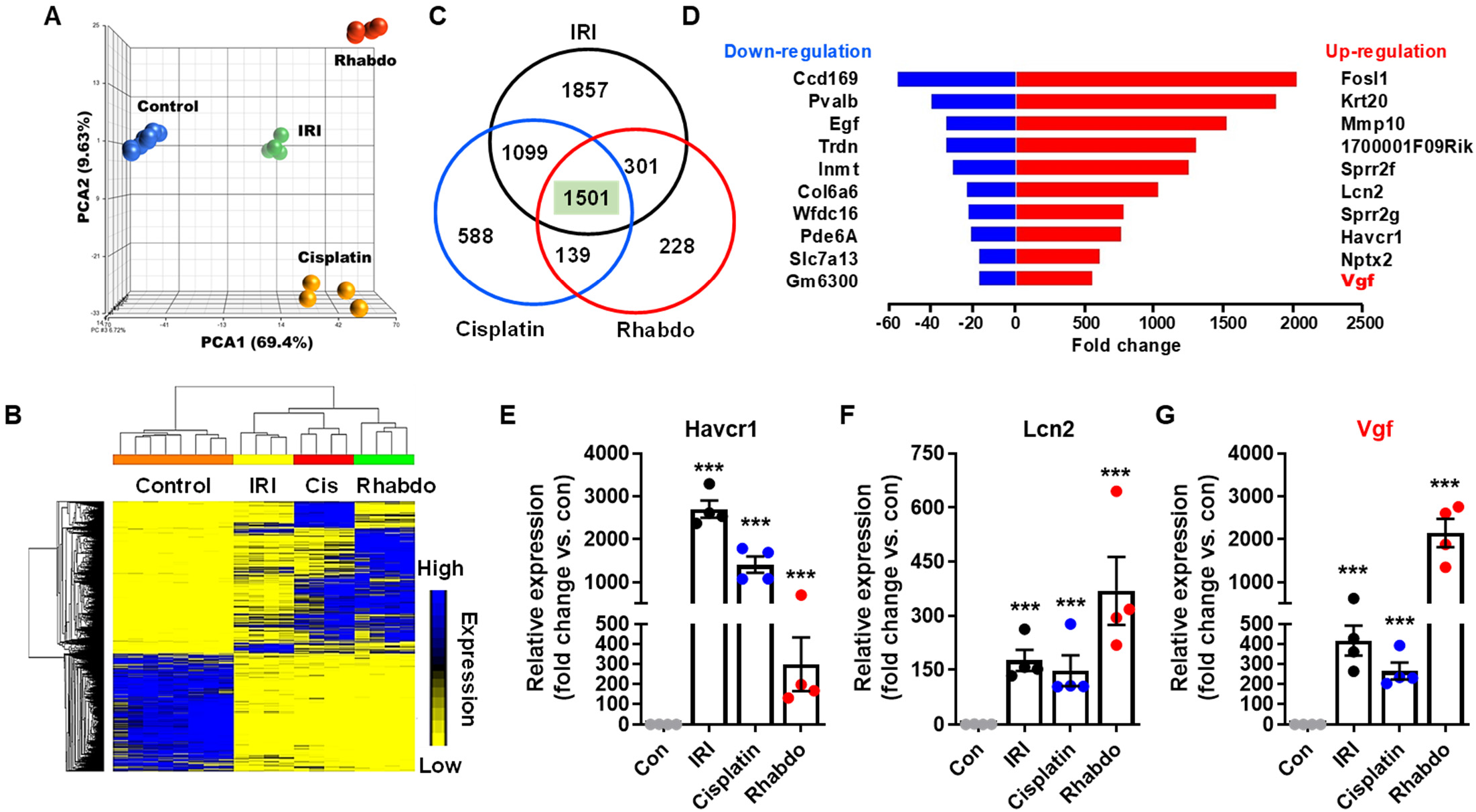
Transcriptome analysis of AKI-associated differentially expressed genes. Renal cortical tissues from control (mock and vehicle treated) mice and mice undergoing AKI (IRI, cisplatin nephrotoxicity, and rhabdomyolysis) were utilized for transcriptome analysis. (**A**) Principal component analysis (PCA) of bulk RNA-Seq data from biological replicates (n=8) of control and AKI (n=4) groups. (**B**) Heatmap of differentially expressed genes during AKI. (**D**) Graphical depiction of top up-and down-regulated genes in three distinct AKI models. (**C**) Venn diagram depicting common differentially expressed genes in the AKI groups. In the three AKI groups, a common set of 1501 genes were found to differentially expressed as compared to control group. (**E-G**) Gene expression analysis of Havcr1, Lcn2, and Vgf genes showed injury induced upregulation. In all the bar graphs, experimental values are presented as mean ± s.e.m. The height of error bar = 1 s.e. and p < 0.05 was indicated as statistically significant. One-way ANOVA followed by Dunnett’s (E-G) was carried out, and statistical significance is indicated by *p < 0.05, **p < 0.01, ***p < 0.001.

To identify previously unexplored genes, we initially focused our attention on the top differentially expressed genes (DEGs) in the AKI conditions. The top common upregulated genes in AKI mice were Fosl1, Krt20, Mmp10, 1700001F09Rik, Sprr2f, Lcn2, Sprr2g, Havcr1, Nptx2, and Vgf (**Fig. 2D**). On the other hand, the top common downregulated genes were Ccdc169, Pvalb, Egf, Trdn, Inmt, Col6a6, Wfdc16, Pde6a, Slc7a13, and Gm6300. The molecular functions of some of these DEGs including the widely studied injury biomarkers Lcn2 and Havcr1 has been explored previously (27,28). We found that similar to Havcr1 and Lcn2 upregulation, 200-2000 fold induction of Vgf gene expression is observed in the three AKI conditions (**Fig. 2E-G**). These results indicated that Vgf is transcriptionally upregulated in response to wide-ranging forms of renal injury.

### Stress-induced Vgf upregulation in RTECs during AKI

Vgf (nonacronymic) was first identified as a nerve growth factor (Ngf) induced gene in a neuroendocrine cell line (18). The Vgf gene encodes a highly conserved precursor polypeptide of 615 (human) and 617 (rat and mice) amino acids. The precursor polypeptide contains several cleavage sites and protease action at these locations results in the generation of a number of peptides, which exert pleotropic biological activities (19), including promotion of pro-survival signaling in an autocrine and paracrine fashion (29,30). While Vgf plays critical roles in neuronal and endocrine tissues, its role in the kidneys remains unknown.

We initially sought to validate the RNAseq data and investigate the cellular origin of Vgf mRNA upregulation. To do so, we utilized a reporter mouse (31) that express membrane-localized green fluorescent protein (GFP) in the tubular epithelial cells (**Fig. 3A**). These mice were challenged with ischemia, cisplatin, and rhabdomyolysis (**Suppl. Fig. 1**) followed by isolation of GFP-positive cells from the kidneys and subsequent examination of Vgf gene expression. We found that Vgf mRNA upregulation occurs in RTECs (GFP-positive cells) early during the development of AKI (**Fig. 3B-D**). A similar increase in Vgf expression was observed when human and murine RTEC cell lines (HK-2 and BUMPT cells) as well as primary murine RTECs were challenged with cisplatin under in vitro conditions (**Fig. 3E**). Based on these results we concluded that Vgf upregulation in RTECs is a common response to stress in vitro and in vivo.

**Figure 3:**
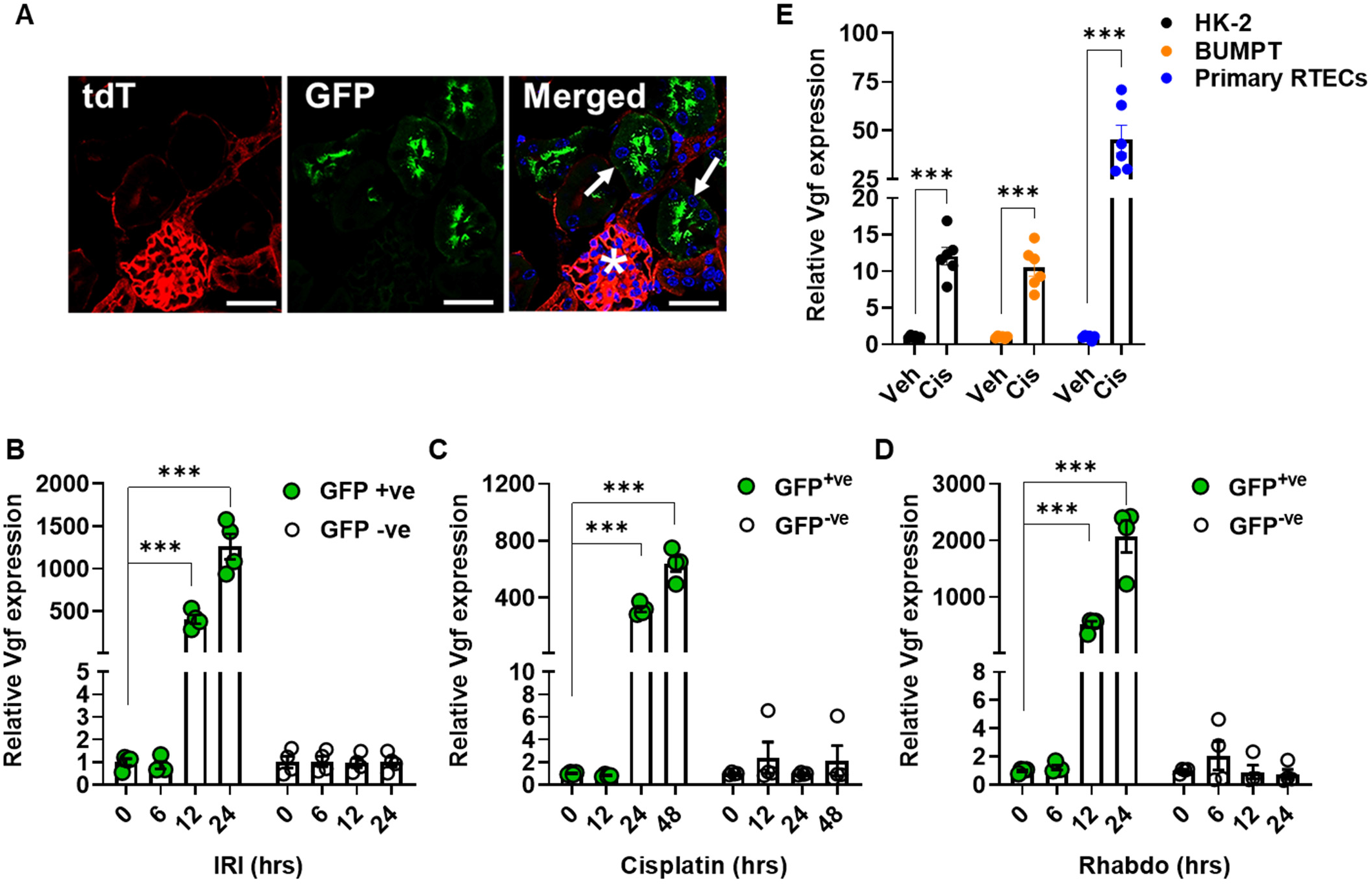
Vgf is upregulated in the tubular epithelial cells during AKI. ROSAmT/mG were crossed with Ggt1-Cre mice to generate transgenic mice that express membrane-localized EGFP in renal tubular epithelial cells, while the other cell types express membrane localized tdTomato. (**A**) Representative image showing EGFP expressing renal tubules (arrows) and tdTomato expression in the glomerulus (asterisks). (**B-D**) Renal Vgf expression was monitored at indicated time-points using qPCR-based analysis. The results demonstrate Vgf gene induction during the early stages of AKI. (**E**) Human (HK-2) and murine (BUMPT) tubular epithelial cell lines as well as murine primary tubular epithelial cells were treated with either vehicle (PBS) or 50 μM cisplatin followed by gene expression analysis at 12 hours. Cisplatin treatment in vitro resulted in Vgf mRNA induction in both the cell lines as well as primary tubular epithelial cells. The graphs (B-E, n = 4-6) are representative of 3 independent experiments. In all the bar graphs, experimental values are presented as mean ± s.e.m. The height of error bar = 1 s.e. and p < 0.05 was indicated as statistically significant. One-way ANOVA followed by Dunnett’s (B-D) or Student’s t-test (E) was carried out, and statistical significance is indicated by *p < 0.05, **p < 0.01, ***p < 0.001. Scale bar (A): 100 μm.

### Vgf gene deletion in renal tubular epithelial cells aggravates AKI

To probe the functional role of Vgf in the pathogenesis of AKI, we examined the effect of Vgf gene ablation on the severity of AKI. To this end, we generated Vgf conditional knockout (Vgf^PT−/−^) mice by crossing the Vgf floxed mice with the Ggt1-Cre mice. In Ggt1-Cre mice, Cre recombinase is expressed in RTECs 7-10 days after birth and as a result Cre-mediated gene ablation is unlikely to influence normal renal development (32). Vgf^PT−/−^mice were indistinguishable from wild-type littermates and normal renal function was not evidently influenced by Vgf deficiency in RTECs (**Suppl. Fig.2**). However, when the control and Vgf^PT−/−^littermates were challenged with ischemia, cisplatin- and rhabdomyolysis, we observed that Vgf gene deletion markedly exacerbates renal injury (**Fig. 4A-I**). Immunoblot analysis of renal tissues confirmed Vgf ablation in the conditional knockout mice (**Fig. 4J**). To further corroborate these results, we cultured primary RTECs from the wild type and Vgf^PT−/−^mice, challenged them with cisplatin and then carried out viability assays. Cell survival and caspase assays (**Fig 4K and Suppl. Fig. 3**) showed that Vgf gene deletion results in increased cisplatin-induced cell death. Thus, we propose that Vgf plays a cytoprotective role in RTECs under stress conditions associated with AKI.

**Figure 4:**
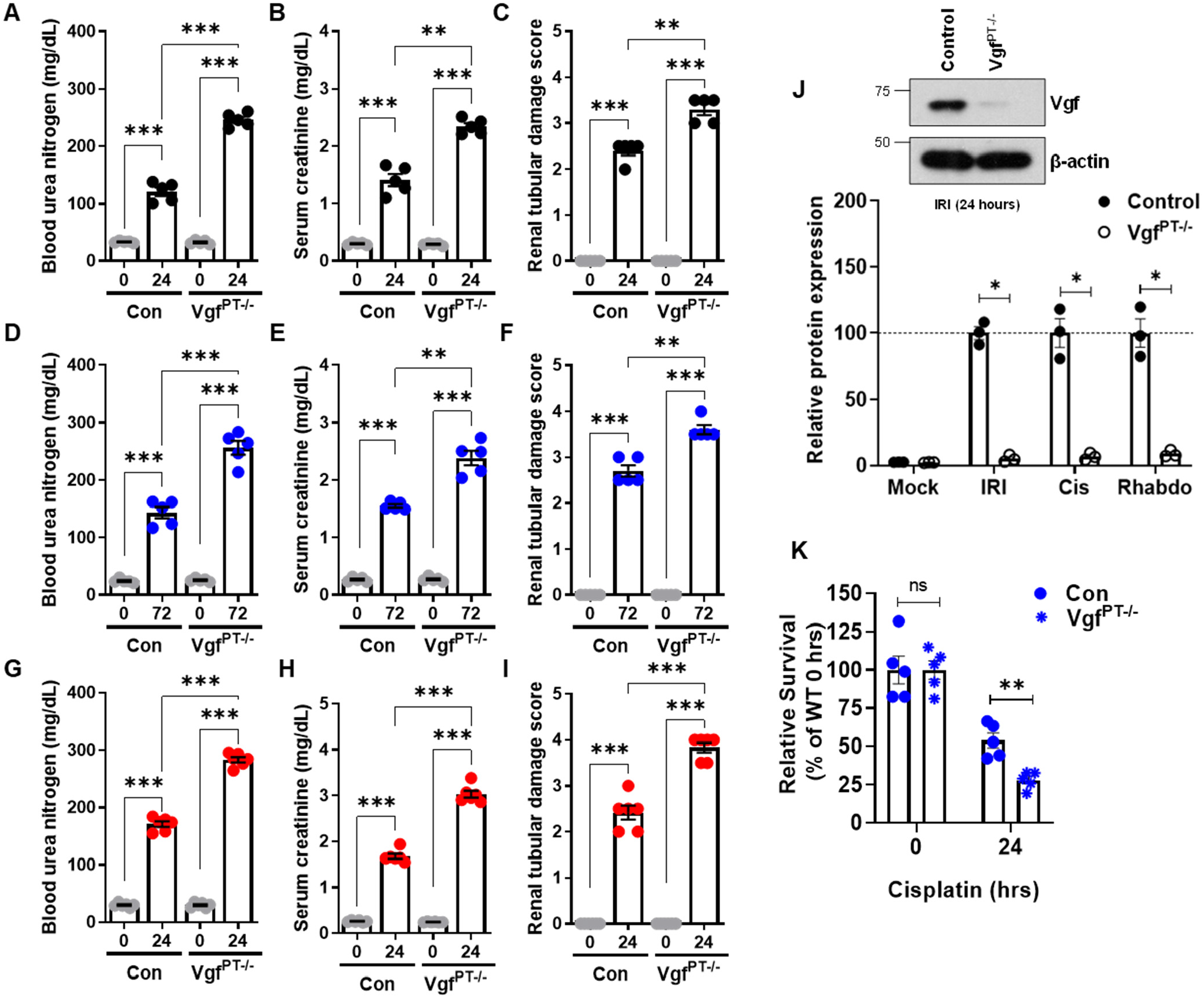
RTEC-specific Vgf gene ablation aggravates AKI. RTEC-specific Vgf knockout mice were generated by crossing Ggt1-Cre mice with Vgf-floxed mice. 8-12 weeks old littermate control and Vgf conditional knockout male mice (indicated by Vgf^PT−/-^) were then challenged with bilateral renal ischemia (30 minutes), cisplatin (30 mg/kg, single intraperitoneal injection) treatment, or glycerol-induce rhabdomyolysis (7.5 ml/kg 50% glycerol in the hind-leg muscles) followed by examination of renal structure and function. Blood urea nitrogen, serum creatinine, and renal histological analysis (H&E) showed that tubular epithelial-specific VGF deficiency results in aggravated renal impairment in the IRI (**A-C**), Cisplatin (**D-F**), and rhabdomyolysis (**G-I**) associated mouse models of AKI. Data presented (A-I) are cumulative of two out of four independent experiments (n = 5) that showed similar results (**J**) Immunoblot analysis of Vgf protein levels was performed using renal cortical tissues from the control and Vgf deficient mice followed by densitometric analysis (normalized to β-actin levels) using ImageJ. The graph depicts relative Vgf protein levels and the upper panel is a representative western blots showing successful gene knockout in the renal tissues. Blots are representative of two independent experiments. (**K**) Primary renal tubular cells isolated from mice with indicated genotypes were treated with 50 μM cisplatin, followed by cell viability assessment using trypan blue staining. Vgf deficiency increased the sensitivity to cisplatin-induced cell death in vitro. Data are presented as individual data points (n = 5 biologically independent samples), from one out of three independent experiments, all producing similar results. In all the bar graphs, experimental values are presented as mean ± s.e.m. The height of error bar = 1 s.e. and p < 0.05 was indicated as statistically significant. One-way ANOVA followed by Tukey’s multiple-comparison test (A-I) was carried out, and statistical significance is indicated by *p < 0.05, **p < 0.01, ***p < 0.001.

### Vgf-associated TLQP-21 peptide protects RTECs under stress conditions

Vgf has several pleiotropic functions in neurons and endocrine cells (18). Notably, several Vgf proteolytic peptides have been identified and are named by the first 4 N-terminal amino acids and their total length (e.g., TLQP-62, TLQP-21, HHPD-41, AQEE-11, and LQEQ-19) (19). These peptides can influence various cellular processes including activation of pro-survival signaling under stress conditions (29,30,33,34). Some of the biological effects of these Vgf-associated peptides are believed to be mediated through binding to extracellular receptors such as the complement-binding protein, gC1qR and complement C3a receptor-1 (C3AR1) (35,36).

We found that TLQP-21 levels were increased in the renal tissues of wild type mice challenged with ischemia, cisplatin and rhabdomyolysis-associated AKI (**Fig. 5A**). Additionally, wild type primary murine RTECs secreted TLQP-21 in the medium when challenged with cisplatin (**Fig. 5B**). Interestingly, tissue distribution studies in mice have shown that intravenously injected TLQP-21 markedly accumulates in the kidney (37). This prompted us to carry out in vivo ‘add-back’ experiments to determine if the TLQP-21 administration can reverse the aggravated renal impairment phenotype observed in the Vgf^PT−/−^mice. To this end, we administered a scrambled peptide (Scr) or TLQP-21 to control and Vgf^PT−/−^mice, 24 and 48 hours after challenging them with cisplatin (**Fig. 5C**). Remarkably, we found that TLQP-21 administration mitigates cisplatin-associated AKI in the Vgf^PT−/−^mice (**Fig. 5D-F**), indicating that loss of TLQP-21 might be partly responsible for the increased sensitivity to renal injury. Complementary studies in primary murine RTECs showed that TLQP-21 treatment can protect Vgf deficient RTECs from cisplatin-induced cell death (**Fig. 5G-I**). These results indicate that the loss of TLQP-21 is partly responsible for the aggravated renal impairment phenotype seen in the Vgf deficient mice.

**Figure 5:**
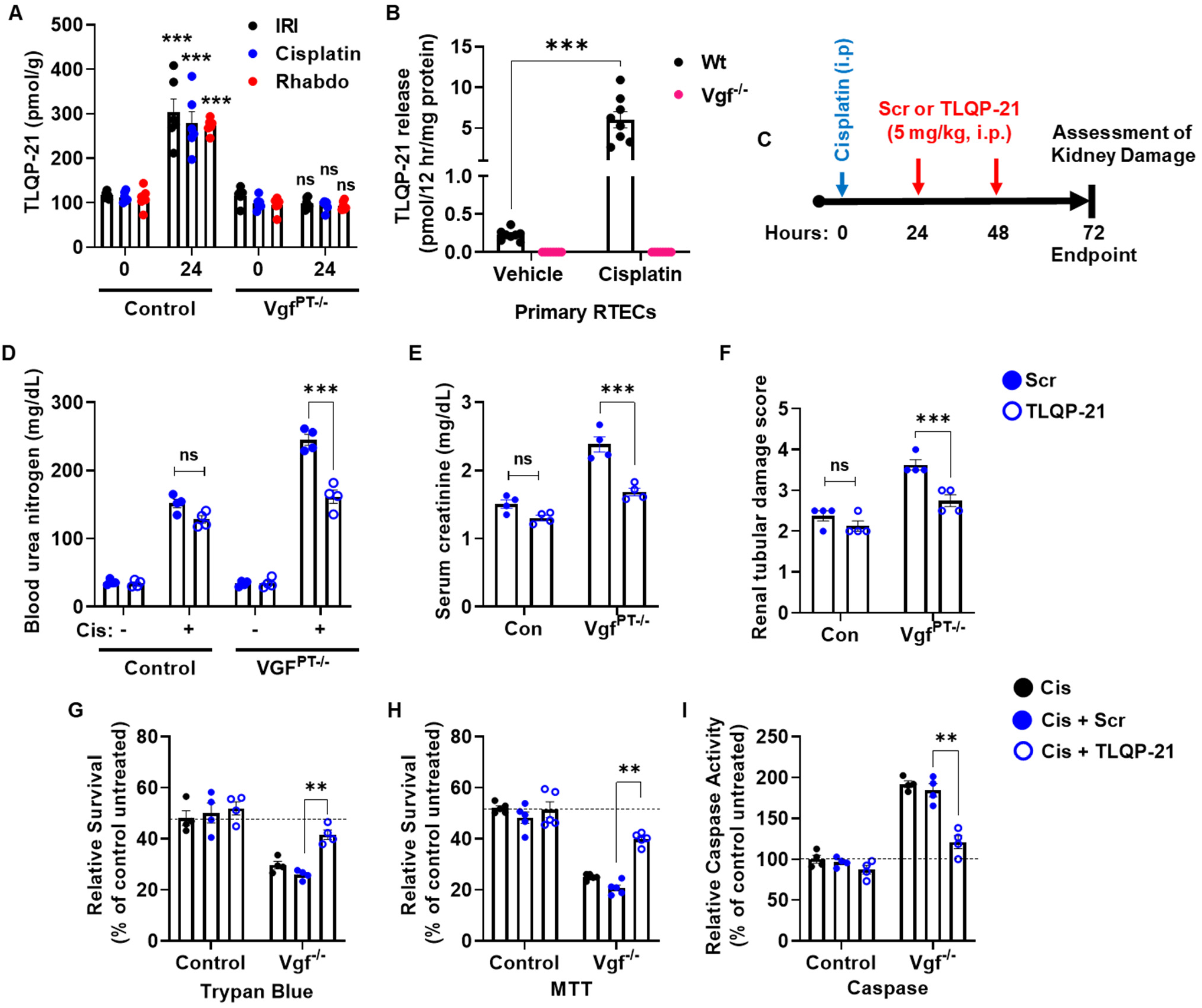
Renal protective effects of Vgf-derived peptide TLQP-21. (**A**) Kidney cortical tissue lysates from mock or vehicle treated mice (-) as well as mice undergoing AKI (IRI-24 hours, Cisplatin-72 hours, and Rhabdomyolysis-24 hours) were used for measurement of TLQP-21 levels using an ELISA based assay. More than 2 fold increase in TLQP-21 levels was observed in the control mice undergoing AKI. * indicates statistical significance as compared to respective mock or vehicle treated (-) group. (**B**) Primary murine renal tubular cells of indicated genotypes were treated with 50 μM cisplatin and levels of TLQP-21 secreted in the media was assessed by an ELISA-based method. (**C**) Littermate control and Vgf^PT−/-^mice were treated with cisplatin (30 mg/kg) on Day 0, followed by intraperitoneal injection of either a scrambled peptide or TLQP-21 (5 mg/kg in normal saline) on day 1 and 2. Renal damage was then examined at day 3. BUN (**D**), serum creatinine (**E**), and histological analysis (**F**) showed that the TLQP-21 administration can partly reverse the aggravated cisplatin nephrotoxicity seen in the Vgf^PT−/-^mice. (**G-I**) Primary murine renal tubular cells of indicated genotypes were treated with 50 μM cisplatin followed by treatment with 25 nM scrambled peptide or TLQP-21 four hours later. At 24 hours cell viability (trypan blue and MTT) and caspase assays were performed, which showed that TLQP-21 can reverse the hyper-sensitive response of Vgf deficient cells to cisplatin treatment. Data (A-B and D-I) are presented as individual data points (n = 4-7 biologically independent samples), from one out of 2-3 independent experiments, all producing similar results. In all the bar graphs, experimental values are presented as mean ± s.e.m. The height of error bar = 1 s.e. and p < 0.05 was indicated as statistically significant. One-way ANOVA followed by Dunnett’s was carried out, and statistical significance is indicated by *p < 0.05, **p < 0.01, ***p < 0.001.

### VGF regulation by Sox9 in the early acute phase of renal injury

Next, we sought to identify the transcriptional mechanisms underlying stress-induced Vgf upregulation in RTECs. While exploring the transcription factor binding sites in the Vgf promoter, we noticed the presence of a putative Sox9 binding site (**Fig. 6A**). Sox9 is upregulated in RTECs in response to injury and is a critical transcriptional regulator of epithelial cell fate during AKI (31,38–41). To test the hypothesis that Sox9 is involved in Vgf upregulation during AKI, we performed promoter-driven luciferase-based reporter assays (**Fig. 6B**) in HEK293 cells, which have low endogenous Sox9 expression. To this end, we used HEK293 cells with stable vector transfection (low Sox9) and Sox9 overexpression (high Sox9) for Vgf promoter driven luciferase reporter assays as described in our recent study (31). We found that Sox9 increases Vgf promoter activity (**Fig. 6C**). Importantly, site-directed mutagenesis of Sox9 binding site within the Vgf promoter suppressed promoter activity. To substantiate these findings, we performed chromatin immunoprecipitation analysis, which confirmed Sox9 binding at the Vgf promoter in vivo (**Fig. 6D**). We next asked if RTEC-specific Sox9 deficiency influences stress responsive Vgf upregulation. Our recent study (31) revealed a protective role of Sox9 during ischemic and nephrotoxic AKI. We also found that RTEC-specific Sox9 gene deletion aggravates rhabdomyolysis-associated AKI **(Suppl. Fig. 4)**. When we carried out gene expression analysis of renal tissues from control and Sox9^PT−/−^mice, we found that stress-induced Vgf upregulation is Sox9 dependent (**Fig. 6E-G**). Taken together, these data indicate that Sox9 controls Vgf gene transcription in RTECs during AKI.

**Figure 6:**
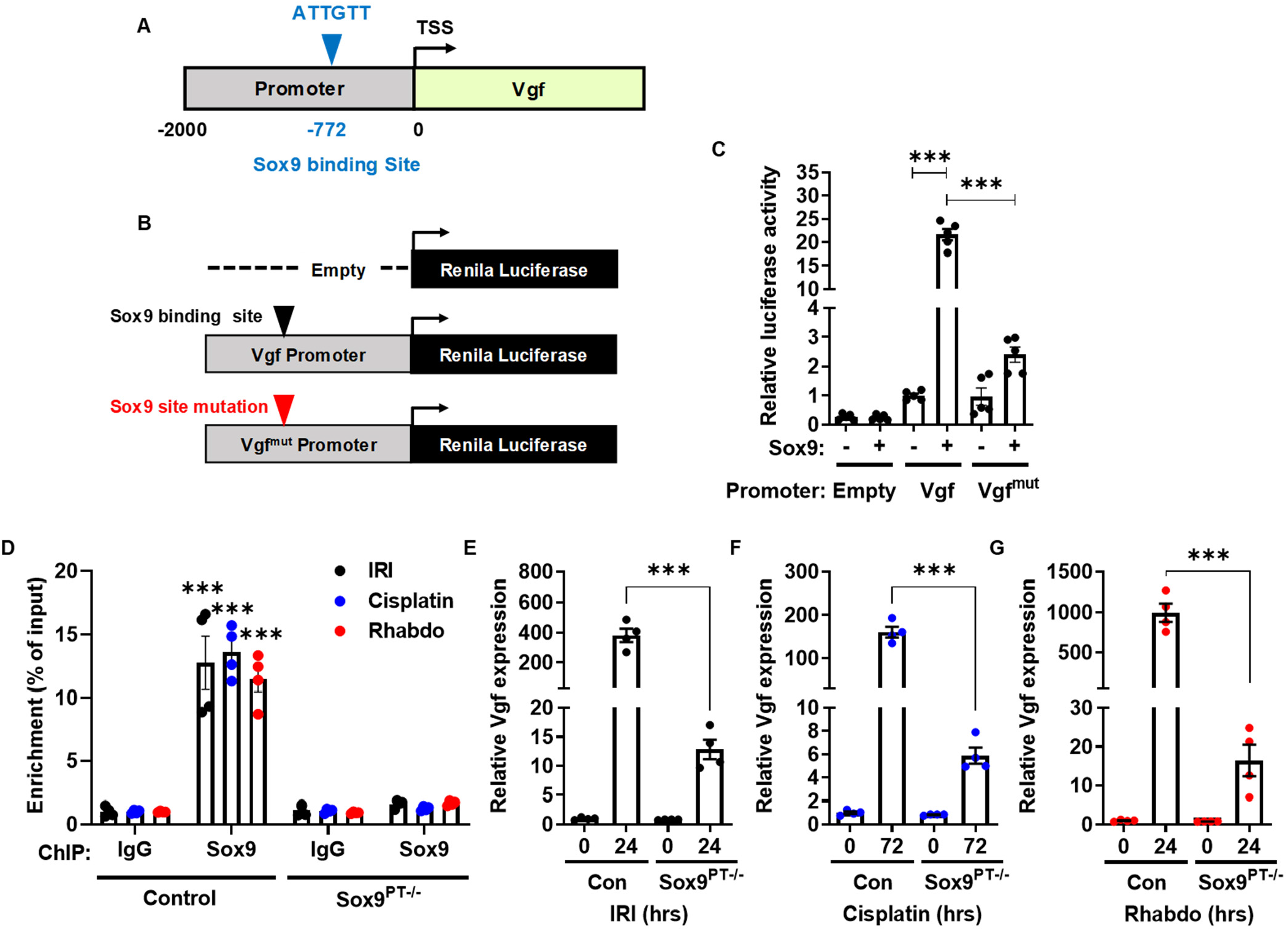
Vgf is a Sox9 target gene. (**A**) Schematic representation of murine Vgf promoter, highlighting the putative Sox9 binding site. (**B-C**) HEK293 cells were stably transfected with empty plasmid or Sox9 construct. The mock transfected cells (-) and Sox9 expressing cells (+) were then used for luciferase based reporter assays. The Vgf promoter sequences (−2000bp from TSS) was cloned in a luciferase reporter construct. Mock (-) and Sox9 (+) expressing cells were transiently co-transfected with reporter renila luciferase constructs (empty, Vgf or Vgf^mut^) and reference cypridina luciferase (normalizing control), followed by measurement of luciferase activity at 24 hours. The normalized luciferase activity of mock (-) group was then compared with the Sox9 (+) group. The results show that the Sox9 can activate transcription from the Vgf promoter. In the Vgf^mut^ construct, the Sox9 binding site was mutated from ATTGTT to AACAAT. (**D**) Sox9 Chromatin immunoprecipitations (ChIP) were carried out from the renal tissues of control and Vgf^PT−/−^mice undergoing Ischemia, cisplatin nephrotoxicity, and rhabdomyolysis associated AKI. Subsequent qPCR analysis using primers specific for the murine Vgf promoter region showed that Sox9 can bind to Vgf promoter in vivo. (**E-G**) q-PCR based gene expression analysis was carried out in the renal tissues from littermate control and Sox9^PT−/−^mice under at indicated time-points after induction of kidney injury. The injury induced Vgf mRNA upregulation was suppressed in the Sox9^PT−/−^mice. Data (C-G) are presented as individual data points (n = 4-5 biologically independent samples), from one out of three independent experiments, all producing similar results. In all the bar graphs, experimental values are presented as mean ± s.e.m. The height of error bar = 1 s.e. and p < 0.05 was indicated as statistically significant. One-way ANOVA followed Tukey’s (C) or Dunnett’s (D-G) multiple-comparison test was carried out, and statistical significance is indicated by *p < 0.05, **p < 0.01, ***p < 0.001.

## Discussion

Here we have mapped the transcriptome changes accompanying the development of ischemic, nephrotoxic, and rhabdomyolysis associated acute kidney injury. We find that these diverse stress conditions trigger transcriptional upregulation of Vgf gene in renal tubular epithelial cells. Importantly, we provide functional evidence that Sox9-mediated Vgf upregulation protects RTECs from cell death and dysfunction linked with AKI. These findings identify Vgf as an essential stress-responsive and protective gene in kidney epithelial cells.

Spatial and temporal changes in gene expression in response to ischemia reperfusion associated kidney injury has been comprehensively explored (39,42). Since multiple etiologies can contribute to the development of AKI, in the current study, we aimed to identify common transcriptional changes that occur in the acute phase of three distinct murine models of AKI. Consistent with previous studies (39,42), we observed that genes such as Sprr2f and Krt20 and well-characterized renal injury biomarkers such as Lcn2 and Havcr1 were significantly upregulated during ischemic, nephrotoxic, and rhabdomyolysis associated acute kidney injury. Furthermore, pathway enrichment analysis revealed that genes linked to cell death and survival, wound healing, small molecule and fatty acid metabolism, and molecular transport were differentially expressed during AKI.

Vgf was among the top upregulated genes in the renal tissues of mice undergoing ischemic, nephrotoxic, and rhabdomyolysis-associated AKI. The Vgf gene is known to be expressed in a subset of cells in the central and peripheral nervous system as well as endocrine cells in the adrenal gland, gastrointestinal tract, and pancreas (18). Within the nervous system, Vgf expression is rapidly induced by neurotrophins, synaptic activity, nerve injury, inflammation, and other stimuli (21). Consistent with its expression in the central and peripheral nervous system, Vgf has been implicated in the regulation of neuroplasticity associated with learning, memory, depression, and chronic pain (21,43,44). Additionally, Vgf plays a critical role in energy homeostasis and metabolism (23,34,45). Mice with germline Vgf deletion are lean, hyper-metabolic, and resistant to diet-, lesion-, and genetically induced obesity and diabetes (20). Interestingly, the role of Vgf in renal physiology and pathology has remained unexplored.

We found that Vgf expression is low in the normal adult kidneys. Moreover, renal epithelial-cell-specific Vgf deficiency did not have any deleterious effect on the normal kidney structure or function and did not alter the overall body weight. Importantly, Vgf expression increased by more than 500 fold in RTECs during ischemic, nephrotoxic, and rhabdomyolysis associated AKI. A previous study (39) also described Vgf gene induction during IRI, however, its functional role in the pathogenesis remained unknown. We find that RTEC-specific Vgf deletion markedly aggravated renal impairment linked with ischemic, nephrotoxic, and rhabdomyolysis-associated AKI. Notably, stress-responsive Vgf upregulation was recapitulated in human and murine cell culture models of cisplatin associated cellular injury. Functional studies also showed that Vgf deficiency sensitizes RTECs to cisplatin-mediated cell death. These studies reveal that Vgf protects RTECs from cell death and dysfunction.

The neuro-endocrine functions attributed to the Vgf gene are dependent on the post-translational processing of Vgf polypeptide into various bioactive peptides, such as TLQP-21, TLQP-62, AQEE-30, LQEQ-19, and NERP2. Among these, TLQP-21 is known to control regulatory processes involved in energy expenditure, lipolysis, glucose-stimulated insulin secretion, gastric acid secretion and pain (34,44,46). We found that along with Vgf mRNA, TLQP-21 levels also increase in the renal tissues during AKI. Strikingly, systemic TLQP-21 administration partly reversed the injury-induced aggravation of renal impairment observed in the RTEC-specific Vgf-deficient mice. These findings suggest that Vgf-derived TLQP-21 plays a protective role during AKI. A critical feature of Vgf peptides is their cell type specific diversity in tissues studied so far and their selective modulation in response to organ or cell type relevant stimuli (19). Future studies are thus necessary to comprehensively profile Vgf derived peptides in renal tissues under normal and stress conditions.

The underlying signaling mechanisms associated with the myriad neuro-endocrine functions of Vgf remains incompletely understood. At least some of the biological functions of Vgf derived peptides are mediated through extracellular receptor binding. Indeed, complement C3a receptor 1 (C3aR1) has been identified as a TLQP-21 receptor on microglia and other cell types (35,47–49). For example, in the adipose tissue, TLQP-21 exerts an anti-obesity effect in diet-induced obese mice through binding to C3aR1 and inducing β-adrenergic receptor expression (50). Furthermore, C1qR, the globular heads of the C1q receptor has been identified as a TLQP-21 receptor in macrophages (36). Interestingly, a previous study has shown that C3aR1 is expressed in RTECs and germline C3aR1 gene ablation provides protection from ischemia-associated AKI (51). It is well established that complement activation within the injured kidneys trigger downstream inflammatory events within the renal parenchyma that exacerbate renal cell dysfunction and cell death (52,53). Future studies with cell type specific conditional knockout mice are required to tease out the possible role of C3aR1, C1qR, or other proteins as receptors of Vgf-derived peptides in the kidney.

How Vgf protects renal epithelial cells under stress conditions in vivo remains unclear. Our studies with cultured primary epithelial cells suggest that stress conditions trigger the induction of Vgf-derived TLQP-21, which might function in an autocrine and or paracrine manner to protect RTECs from cisplatin-associated cell death. However, these epithelial cell culture models of injury do not completely recapitulate the in vivo pathophysiological complexities of AKI, particularly the involvement of other cell types such as immune cells (54–56). Therefore, our studies do not rule out the possibility that Vgf and TLQP-21 might influence renal injury through crosstalk between epithelial and immune cells. Interestingly, in peripheral neurons, inflammatory conditions can cause Vgf upregulation and Vgf can in turn functionally regulate inflammatory processes (57). Altogether, our study provides strong evidence for the protective role of RTEC-derived Vgf in AKI, however, the further unravelling of this pathway will require the identification of underlying receptors, modulated cell types and intracellular signaling pathways.

While the downstream pathways remain unclear, we propose that Sox9 is the critical transcriptional regulator of stress-induced Vgf upregulation in RTECs. Several lines of evidence suggest that Sox9 directly binds to the Vgf promoter and promotes the transcriptional up-regulation of Vgf gene. Firstly, injury-induced Vgf upregulation was significantly suppressed in renal tissues of RTEC-specific conditional *Sox9* knockout mice. Secondly, chromatin immunoprecipitation studies showed Sox9 enrichment at the Vgf promoter site in vivo. Thirdly, luciferase reporter assays confirmed Sox9 mediated transactivation of Vgf promoter, which was suppressed by mutations in the Sox9 binding site. These results establish Vgf as a bona fide Sox9 target gene in RTECs.

In the current study, we find that Vgf deficiency exacerbates AKI, a phenotype that is similar to the RTEC-specific Sox9 deficient mice (31). However, Sox9 is a crucial transcriptional regulator of not only the early pathogenic phase (31), but also the later recovery phase of AKI (38,40). Sox9 expressing RTECs are involved in the repair and regeneration processes post-AKI (38,40). Based on our findings that Vgf is a downstream Sox9 target gene, it will be interesting to examine if Vgf contributes to repair and regeneration. It would also be interesting to examine if systemic TLQP-21 administration can accelerate the recovery and repair processes post-AKI. Collectively, our study has revealed Vgf as an essential Sox9 target gene that protects RTECs under stress conditions associated with acute kidney injury.

## Experimental procedures

### Cell culture and reagents

Boston University mouse proximal tubule cells (BUMPT, clone 306, generated by Drs. Wilfred Lieberthal and John Schwartz, Boston University School of Medicine, Boston, MA, were obtained from Dr. Zheng Dong, Augusta University, Augusta, GA) were grown at 37 °C in Dulbecco’s modified Eagle’s medium with 10% fetal bovine serum. The human renal tubular cell line, HK-2 cells (ATCC, CRL-2190) were grown in keratinocyte media (K-SFM) according to the provider’s instructions. HEK293 cells stably transfected with empty vector (pCMV6) or Sox9 expression vector have been described in our previous study (31) and were grown at 37 °C in Dulbecco’s modified Eagle’s medium with 10% fetal bovine serum. Cisplatin, glycerol, TLQP-21 (murine), and other reagents were obtained from Sigma-Aldrich.

### Primary murine tubular cell culture

Murine renal cortical tissues were minced and digested with 0.75 mg/ml collagenase IV (Thermo-Fisher Scientific). Cells were centrifuged at 2000 g for 10 min in DMEM/F-12 medium with 32% Percoll (Amersham). After two washes with serum-free media, the cells were plated in collagen-coated dishes and cultured in DMEM/F-12 medium supplemented with 5 μg/ml transferrin, 5 μg/ml insulin, 0.05 μM hydrocortisone, and 50 μM vitamin C (Sigma-Aldrich). Fresh media was supplemented every alternate day, and after 5–7 days of growth, the isolated proximal tubular cells were trypsinized and re-plated at 1 × 10^5^ cells per well in 24-well plates. To induce cell death, primary RTECs were incubated with 50 μM cisplatin (Sigma-Aldrich) in fresh culture medium for 24 h, followed by viability and caspase assays.

### Cell viability and caspase assays

Trypan blue staining and MTT assay was used to determine cellular viability as reported in our previous study (31). BUMPT, HK-2 cells or RTECs were seeded in 6-well, 24-well, or 96-well plates, followed by cisplatin treatment for 24–48 h. At the end of the incubation period, cells from 6-well plates were harvested, followed by trypan blue staining and manual cell counting with a hemocytometer and/or by using Countess Automated Cell Counter (Thermo Fisher); translucent cells were considered as viable and blue-stained cells were counted as dead. Cellular viability was calculated by dividing the number of viable cells by the total cell number and each sample was done in triplicate. For MTT assays, after cisplatin treatment, 10 μL of MTT reagent (5 mg/mL MTT in PBS) was added to each well, and plates were incubated at 37 °C with 5% CO2 for 4 h, followed by addition of 100 μl of acidified isopropanol (Sigma-Aldrich) and measurement of absorbance at 590 nm. The half-maximal inhibitory concentration (IC50) was calculated by nonlinear regression analysis using GraphPad Prism.

For caspase assays (58), RTECs were lysed in a buffer containing 1% Triton X-100, and 10 μg of protein from cell lysates was added to an enzymatic assay buffer containing 50 μM DEVD-AFC for 60 min at 37 °C. Fluorescence at excitation 360 nm/emission 535 nm was measured, and free AFC was used to plot a standard curve. Subsequently, the standard curve was used to convert the fluorescence reading from the enzymatic reaction into the nM AFC liberated per mg protein per hour as a measure of caspase activity.

### Mice strains and breeding

Mice were housed in a temperature-controlled environment with a 12 hour light cycle and were given a standard diet and water ad libitum. All animal experiments were carried out in accordance with the animal use protocol approved by the Institutional Animal Care and Use Committee of the Ohio State University. C57BL/6J mice, Vgf floxed mice, Sox9 floxed mice and Ggt1-Cre transgenic mice (stock numbers 000664, 030571, 013106 and 012841, respectively) were obtained from Jackson Laboratories. Vgf floxed mice and Sox9 floxed mice were bred with Ggt1-Cre transgenic mice to generate conditional gene knockout mice in renal tubular epithelial cells. These transgenic mice express Cre recombinase in the renal tubular epithelial cells beginning at age 1-2 weeks. mT/mG mice that express membrane-targeted, two-color fluorescent Cre-reporter allele were obtained from Jackson Laboratories (stock no. 007676). The mT/mG mice were bred with Ggt1-Cre strain.as reported previously (31). For all mouse colonies, the pups were ear tagged and genotyped at 3 weeks of age. Offspring were genotyped by standard PCR-based methods. Primers used for amplification were Vgf36100 (5′-TCC TCC CTC TCA GTG TTT GC-3′) and Vgf36101 (5′-GGA CTC GCA CAA ACC ACA C-3’) and yield a 313-bp product for the *Vgf* floxed allele, and a 194-bp product for the *Vgf* wild-type allele. Sox9-11576 (5’-AGA CTC TGG GCA AGC TCT GG-3’) and Sox9-11577 (5’-GTC ATA TTC ACG CCC CCA TT-3’) were used for amplification and yield a 300-bp product for *Sox9* floxed allele and a 250-bp product for the *Sox9* wild-type allele. Primers for Ggt1-Cre are Cre5’ (AGG TGT AGA GAA GGC ACT TAG C), Cre3’ (CTA ATC GCC ATC TTC CAG CAG G) and produce a 405-bp product. PCR products were analyzed by electrophoresis using 1.5% agarose gels.

### Animal models of acute Kidney injury

We carried out all the studies presented here in age-matched male mice at 8-12 weeks of age using methods described in our recent studies (31,59,60). In all the studies with conditional Vgf and Sox9 knockout mice, we used male littermates from mice bred in-house. Experiments were carried out in a blinded fashion where the investigators assessing, measuring or quantifying experimental outcomes were blinded to the genotype or treatment of the mice. For ischemia-reperfusion experiments, mice were anesthetized by isoflurane and placed on a surgical platform where the body temperature was monitored throughout the procedure. The skin was disinfected, kidneys were exposed and bilateral renal pedicles were clamped for 30 minutes. Consequently, the clamps were removed to initiate reperfusion followed by suturing to close the muscle and skin around the incision. To compensate for the fluid loss, 0.5 ml warm sterile saline was administered via intraperitoneal injection. Blood was collected on day 1 via cardiac puncture after carbon dioxide asphyxiation. Renal tissues were collected and processed for RNA-*seq*, qPCR, and histological analysis as described previously. For nephrotoxicity experiments, cisplatin (30 mg/kg) was administered by *i.p.* injection as described previously. After cisplatin injection, blood was collected on days 0-3 by submandibular vein bleed or on day 3 via cardiac puncture after carbon dioxide asphyxiation. Renal tissues were collected and processed for RNA-*seq*, qPCR, and histological analysis. To induce rhabdomyolysis, 8-12 weeks old male mice were injected with 7.5 ml/kg 50% glycerol intramuscularly to the two hind-legs or injected with saline as a control, followed by tissue collection at 24 hours and RNA-seq, qPCR, and histological analysis

### Assessment of renal damage

Renal damage was assessed by serum analysis (blood urea nitrogen and creatinine) and histological examination (H&E staining). Mouse blood samples were collected at indicated time-points, followed by blood urea nitrogen and creatinine measurement by QuantiChromTM Urea Assay Kit (DIUR-100) and Creatinine Colorimetric Assay Kit (Cayman Chemical). For histological analysis, mouse kidneys were harvested and embedded in paraffin at indicated time-points before and after AKI induction. Tissue sections (5 μm) were stained with hematoxylin and eosin by standard methods (60). Histopathologic scoring was conducted by in a blinded fashion by examining ten consecutive 100x fields per section from at least three mice per group. Tubular damage was scored by calculation of the percentage of tubules that showed dilation, epithelium flattening, cast formation, loss of brush border and nuclei, and denudation of the basement membrane. The degree of tissue damage was scored based on the percentage of damaged tubules as previously described: 0: no damage; 1: <25%; 2: 25–50%; 3: 50–75%; 4: >75%.

### RNA-seq

Total RNA was isolated from harvested renal cortical tissues using the RNeasy Plus Mini Kit (Qiagen, Germantown, MD, USA) according to the manufacturer’s protocols. Total RNA samples used for library construction and sequencing (Quick Biology, Pasadena, CA). RNA integrity, quality and purity were analyzed by Agilent 2100 Bioanalyzer. Libraries for RNA-Seq were prepared with KAPA Stranded mRNA-Seq Kit (KAPA Biosystems, Wilmington, MA) according to the manufacturer’s protocols. The workflow consisted of mRNA enrichment using bead capture for poly-A selection, cDNA generation, end repair to generate blunt ends, A-tailing, adaptor ligation and PCR amplification of library fragments. Final Library size distribution was determined by using an Agilent 2100 Bioanalyzer using the High-Sensitivity DNA Kit and its quantity was analyzed by Life Technologies Qubit 3.0 Fluorometer. Libraries were pooled and sequenced on the Illumina HiSeq 4000 platform to obtain 150-bp paired-end reads, 20 million reads (10 million reads pairs) per sample.

### Bioinformatics analysis

Sequencing data quality checks were performed by using FastQC followed by read alignments using Bowtie2 version 2.1.0 with alignment to the mouse Ensembl genome (GRCm38/mm10). The overall mapping rate of more than 80% and rRNA percentage less than 5% was considered as good quality mapping data. The reads were first mapped to the latest UCSC transcript set using Bowtie2 (version 2.1.0) and the gene expression level was estimated using RSEM v1.2.15. Differentially expressed genes were called for each time point with Bioconductor edgeR. TMM (trimmed mean of M-values) method in edgeR package was used to normalize the gene expression results. For all the analyses, we only kept genes with (a) FDR-transformed P values below 0.05, (b) fold change of at least 1.5, and (c) TMM above 1 in three distinct AKI and/or control samples. These values of fold change and TMM thresholds were chosen to enable experimental validation of our differential-expression calls. We used the fold changes calculated by edgeR to create a pre-ranked gene-list. Each list of differentially expressed genes derived from the different comparisons were subjected to functional and biochemical pathway analysis using the Gene Ontology (GO) and KEGG, Reactome pathway databases. Goseq was used to perform the GO enrichment analysis and Kobas was used to perform the pathway analysis. The RNA-Seq data have been deposited in the Gene Expression Omnibus (GSE153625).

### qPCR analysis

One microgram of total RNA from renal cortical tissues or cultured RTECs was reversed transcribed using RevertAid First Strand cDNA Synthesis Kit (Thermo-Fisher Scientific) and qRT-PCR was run in QuantStudio 7 Flex Real-Time PCR System (Thermo-Fisher Scientific) using SYBR Green Master Mix and gene-specific primers. The expression levels of the samples were determined by the comparative CT (ΔΔ^CT^) method. β-actin was used as the internal control. For gene expression analysis in RTECs in vivo, anti-GFP antibody and MACS columns (Miltenyi Biotech) were used to isolate GFP-positive tubular epithelial cells form the kidneys of reporter mice with membrane localized EGFP as reported previously (31).

### Immunoblot analysis and ELISA

Whole-cell lysates from renal cortical tissues were prepared using modified RIPA buffer (20 mM Tris-HCl (pH 7.5), 150 mM NaCl, 1 mM Na2EDTA, 1 mM EGTA, 1% NP-40, 2.5 mM sodium pyrophosphate, 1 mM beta-glycerophosphate, protease, and phosphatase inhibitors) supplemented with 1% SDS. Invitrogen Bis-tris gradient midi-gels were used for western blot analysis, followed by detection by ECL reagent (Cell Signaling). Primary antibodies used for western blot analysis were from Santa Cruz Biotech: Vgf (sc-365397), and β-actin (47778) and were used at 1:1,000 dilution. Secondary antibodies were from Jackson Immuno-research and were used at 1:2,000 dilutions. Uncropped images of western blots are shown in **Supplementary Figure 5**. Densitometric analysis was carried out using Image J, and the signals of target protein was normalized to actin levels in the same samples.

For measurement of TLQP-21 secretion in the media, primary murine RTECs were treated with vehicle or 50 μM cisplatin followed by media collection after 12 hours. Secreted TLQP-21 was assayed from the media using a mice TLQP-21-specific ELISA (Peninsula Laboratories International, S-1477) and normalized to cellular protein levels (BCA assay, Thermo Fischer). Similar methodology was used to measure TLQP-21 levels in cellular lysates. Vgf knockout samples were used as negative controls in all the experiments and TLQP-21 levels were undetected in the Vgf deficient cells and tissues.

### TLQP-21 addback experiments

Murine purified TLQP-21 (Sigma, T1581) and a scrambled control peptide (GenScript) were used for addback experiments. For in vivo experiments, littermate control and Vgf deficient male mice (8-12 weeks) were injected with cisplatin (30 mg/kg, intraperitoneal). Twelve hours later, TLQP-21 or control scrambled peptide was administered by intraperitoneal injection (4.5 mg/kg). Renal tissues and blood was collected at 72 hours post cisplatin injection, followed by assessment of renal damage. For in vitro experiments, primary RTECs were sequentially treated with 50 μM cisplatin (or vehicle), followed by TLQP21 or Scrambled peptide (25 nM) treatment four hours later and assessment of cellular viability at 24 or 48 hours.

### Promoter Luciferase Assay

HEK293 cells were stably transfected with either empty vector (pCMV6) or Sox9 expression vector (Origene). These cells were then utilized for promoter luciferase reporter assays using methods reported in our recent studies (31,61). Briefly, (5 × 10^3^) were plated overnight on white poly-l-lysine-coated 96-well plates, followed by transient transfection with either promoter constructs (Switchgear Genomics, encoding approximately 2 kb sequence upstream of transcription start site of Vgf) or empty promoter construct at 30 ng in combination with the Cypridina TK control construct (Switchgear Genomics) at 1 ng, according to the manufacturer’s protocol (Switchgear Genomics, Lightswitch Dual Assay kit, DA010). The promoter construct encodes a Renilla luminescent reporter gene, called RenSP, while the transfection and normalization vector encodes a Cypridina luciferase. The Renilla luciferase activity was normalized to the Cypridina luciferase activity.

### Site directed mutagenesis

The QuikChange II XL Site-Directed Mutagenesis Kit (Agilent) was utilized to generate Vgf promoter mutants, according to previously described methods (62). The QuikChange primer design program was used to design mutagenesis primers and primers were synthesized by Integrated DNA Technologies. Mutant constructs were sequenced to confirm successful mutagenesis. The primers used for Sox9 binding site mutagenesis (ATTGTT to AAACAT) in the Vgf promoter reporter construct were 5’-TGTTCCCTGGTCCATGTTTAAGTTCAAGCCGACAGCATCACCCAG-3’ and 5’-CTGGGTGATGCTGTCGGCTTGAACTTAAACATGGACCAGGGAACA-3’.

### Chromatin immunoprecipitation (ChIP)

ChIP assays were performed using the Pierce Magnetic ChIP Kit according to the manufacturer’s instructions and our previous studies (31,61). Briefly, cross-linking with 1% formaldehyde was carried out in renal tissues, followed by quenching with glycine, harvesting, and DNA fragmentation by sonication. Tissue lysates were precleared for 2 h with Protein A + G magnetic beads (EMD Millipore). Precleared lysates were then incubated with 5 μg of anti-Sox9 antibody (Abcam, ab3697) overnight at 4 °C, followed by addition of Protein A + G magnetic beads and incubation for 4 h at 4 °C. Finally, the beads were collected, repeatedly washed and the protein–DNA complexes were eluted, cross-links were reversed and the DNA was purified. Standard qPCR analysis was then carried out using the following primers spanning the Vgf promoter: 5’-TCCCAGGCTGATGTGAACTT-3’ and 5’-TCACCAGGCATGCCCATAAG-3’.

### Statistical Analysis

Data in all the graphs are presented as mean with s.e.m, unless stated otherwise. Statistical calculations were carried our using GraphPad Prism. p<0.05 was considered as statistically significant. To calculate statistical significance between two groups, two-tailed unpaired Student’s t test was performed. One-way ANOVA followed by Tukey’s or Dunnett’s multiple-comparison test was used for comparisons among three or more groups. No sample outliers were excluded.

## Data Availability

The RNA-Seq data have been deposited in the Gene Expression Omnibus (GSE153625). And the rest of data are contained within the manuscript.

## Acknowledgments

We thank Simarjot Pabla and Sithara Raju Ponny (University of Massachusetts) for assistance with RNAseq data analysis and submission. We also thank Eric Liao and Sophia Wang (Quick Biology, Pasadena, CA) for assistance with transcriptome analysis. We thank Dr. Zheng Dong (Augusta University) for providing the BUMPT cell line, which was originally obtained from Drs. Wilfred Lieberthal and John Schwartz, Boston University School of Medicine, Boston, MA. We thank all members of our laboratories for helpful discussions of this study and critical reading of the manuscript.

## Author contributions

J.Y.K and N.S.P designed research; J.Y.K, Y.B., L.A.J, A.B., F.A., and N.S.P performed research; J.Y.K., M.G., S.V.P, M.S., A.B., and N.S.P analyzed data and J.Y.K and N.S.P wrote the paper.

## Funding and additional information

This study was supported by funds from the Ohio State University Comprehensive Cancer Center and Scientist Development Grant from the American Heart Association (17SDG33440070).

## Abbreviations

AKI: Acute kidney injury
RTEC: renal tubular epithelial cells
BUN: blood urea nitrogen
ChIP: chromatin immunoprecipitation
Sox9: SRY-Box transcription factor 9.

---

**Supplementary Information Figure S1.**
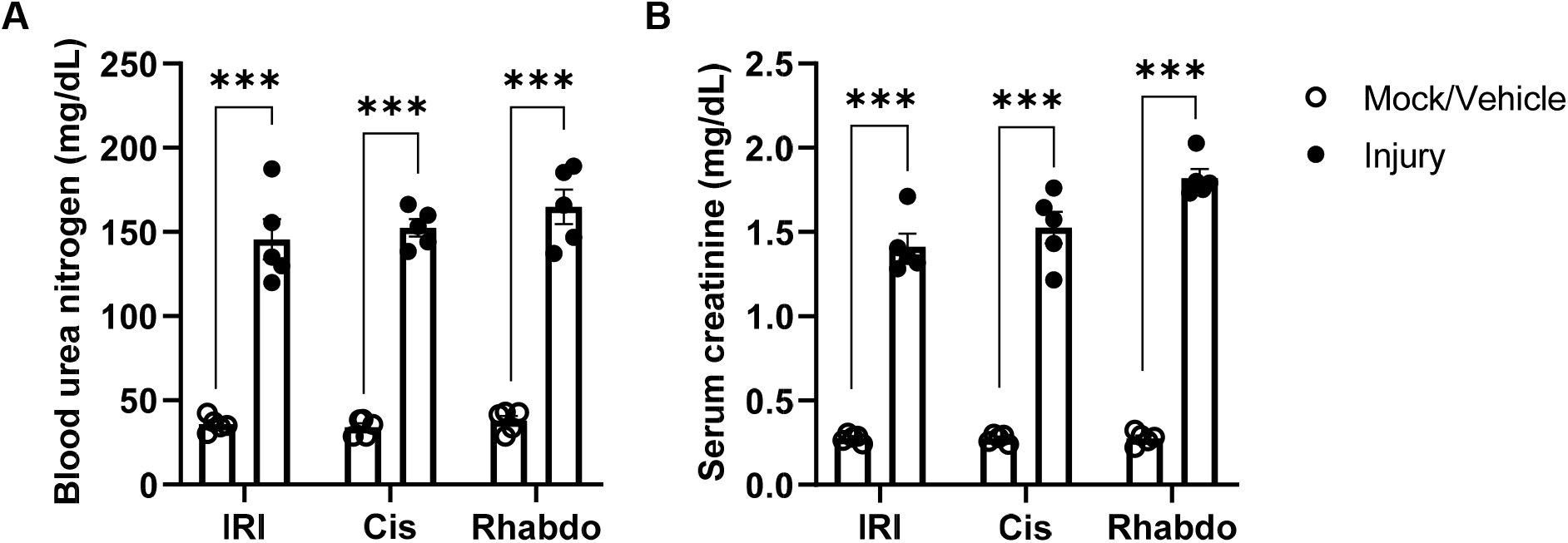
Characterization of the RTEC-specific EGFP expressing reporter mice. Ggt1-Cre mice were crossed with ROSAmT/mG mice to generate transgenic mice that express membrane-localized EGFP in RTECs. 8-12 weeks male mice were then challenged with bilateral renal ischemia (30 minutes), cisplatin (30 mg/kg, single intraperitoneal injection) treatment, or glycerol-induced rhabdomyolysis (7.5 ml/kg 50% glycerol in the hind-leg muscles) followed by examination of renal structure and function. The mock/vehicle groups represent respective control groups (with no injury). (**A-B**) Representative graphs depicting injury-induced increase in blood urea nitrogen and serum creatinine levels (IRI and Rhabdo at 24 hours and Cisplatin at 72 hours). The graphs (n = 5) are representative of three independent experiments. In all the bar graphs, experimental values are presented as mean ± s.e.m. The height of error bar = 1 s.e. and p < 0.05 was indicated as statistically significant. Student’s t test was carried out, and statistical significance is indicated by *p < 0.05, **p < 0.01, ***p < 0.001.

**Supplementary Information Figure S2.**
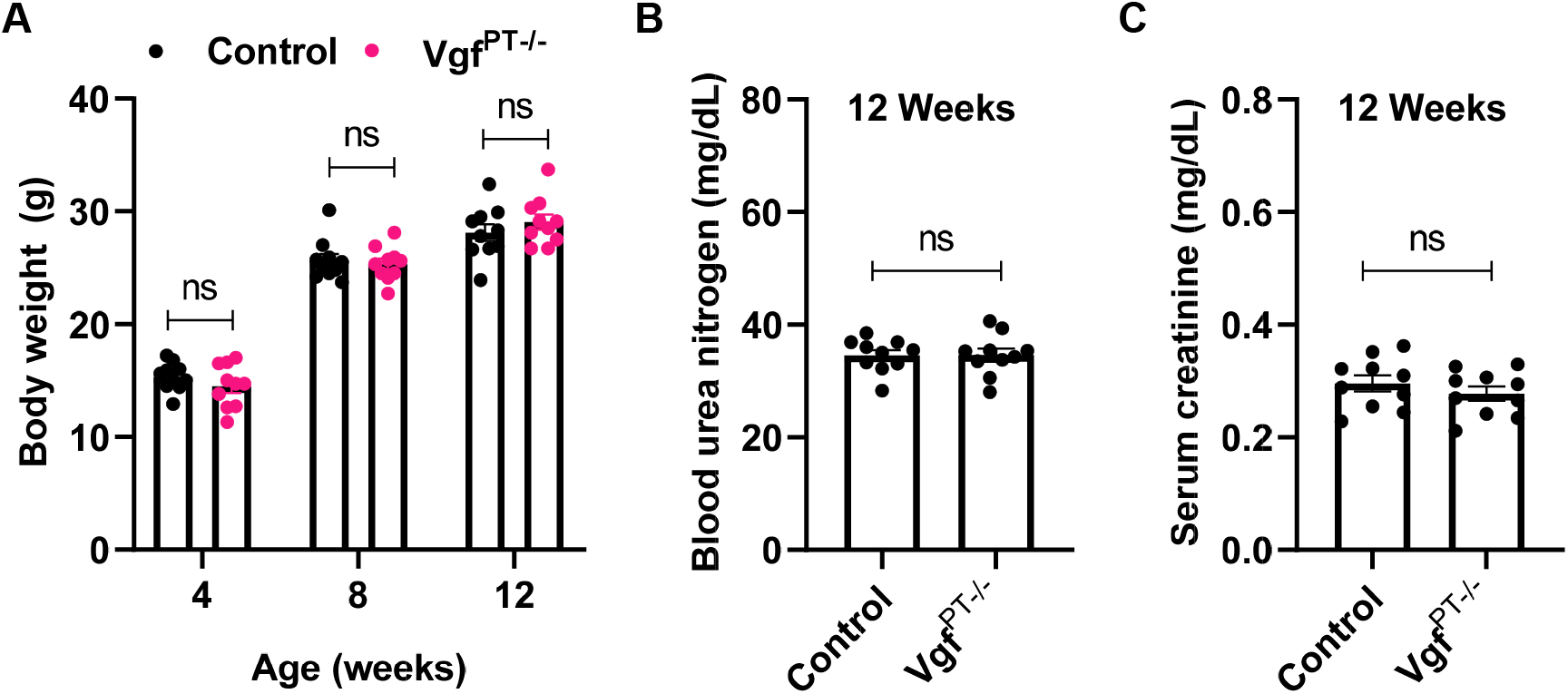
Characterization of the RTEC-specific Vgf deficient mice. Ggt1-Cre mice were crossed with Vgf-floxed mice to generate RTEC-specific Vgf deficient mice. (**A**) Body weight measurements showed no differences between the control and Vgf deficient mice up to 12 weeks of age. (**B-C**) Renal function (BUN and Creatinine) was examined in littermates with indicated genotypes at 12 weeks of age under baseline conditions. These results show that RTEC-specific Vgf knockout does not affect kidney function under normal conditions. Data are presented as individual data points (n = 10), from a single long-term experiment. In all the bar graphs, experimental values are presented as mean ± s.e.m. The height of error bar = 1 s.e. and p < 0.05 was indicated as statistically significant. Student’s t test was carried out, and statistical significance is indicated by *p < 0.05, **p < 0.01, ***p < 0.001, and ns= not significant.

**Supplementary Information Figure S3.**
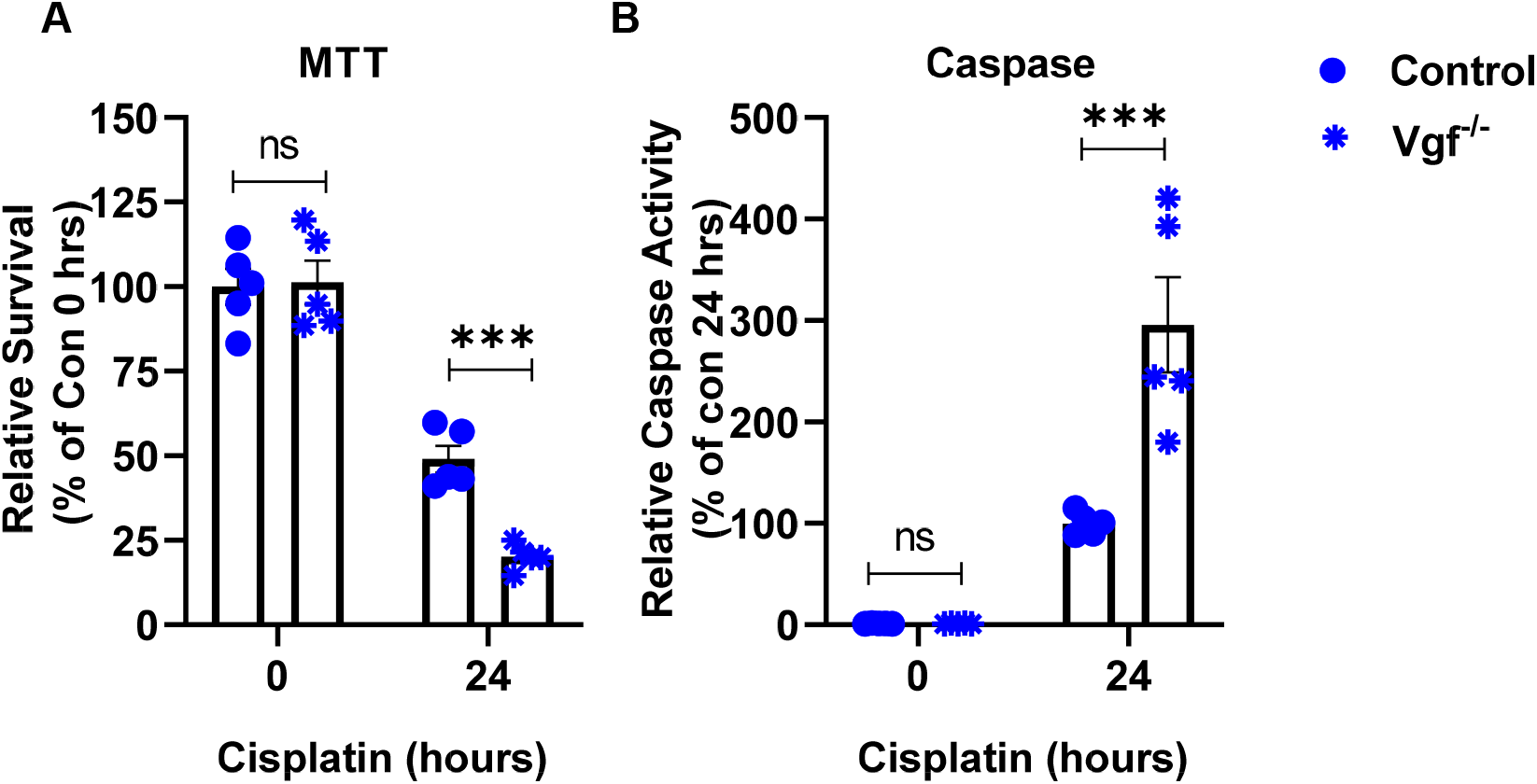
Vgf deficiency sensitizes RTECs to cisplatin-induced cell death. **(A-B)** Primary RTECs from mice with indicated genotypes were treated with 50 μM cisplatin, followed by cell viability assessment using MTT assay and measurement of caspase activity in cellular lysates. Data are presented as individual data points (n = 5 biologically independent samples), from one out of two independent experiments, all producing similar results. In all the bar graphs, experimental values are presented as mean ± s.e.m. The height of error bar = 1 s.e. and p < 0.05 was indicated as statistically significant. One-way ANOVA followed by Dunnett’s was carried out, and statistical significance is indicated by *p < 0.05, **p < 0.01, ***p < 0.001

**Supplementary Information Figure S4.**
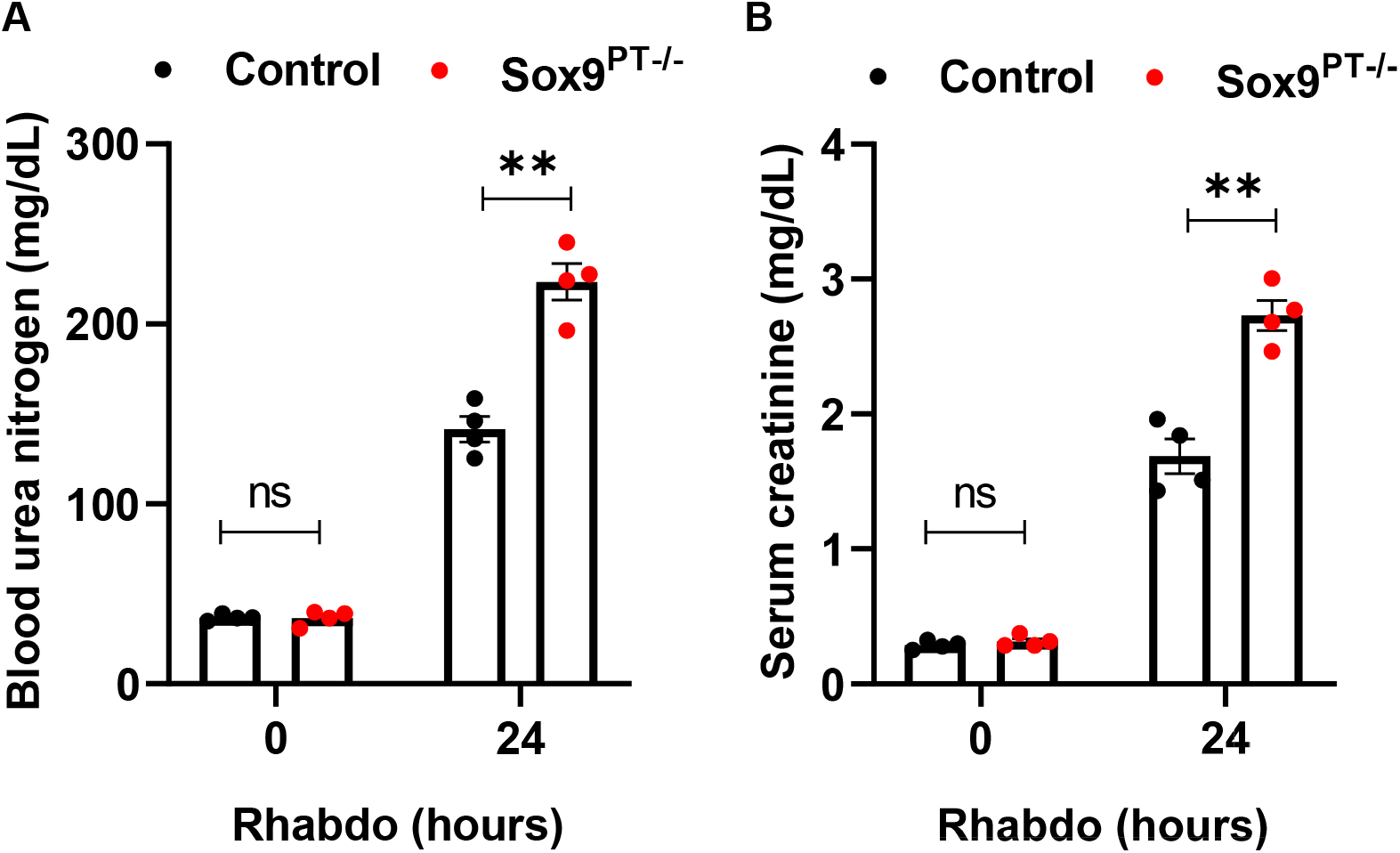
Sox9 plays a protective role during Rhabdomyolysis-associated AKI. To generate mice with renal tubule-specific Sox9 knockout, Ggt1-Cre mice were crossed with Sox9-floxed mice. Control and Sox9^PT−/−^male litermates (8-12 weeks age) were challenged with glycerol-induced rhabdomyolysis (7.5 ml/kg 50% glycerol in the hind-leg muscles) followed by examination of renal impairment. (**A-B**) Blood urea nitrogen and serum creatinine measurements showed that RTEC-specific Sox9 deficiency exacerbates rhabdomyolysis-associated AKI. Data are presented as individual data points (n = 4), from one out of 3-4 independent experiments. In all the bar graphs, experimental values are presented as mean ± s.e.m. The height of error bar = 1 s.e. and p < 0.05 was indicated as statistically significant. One-way ANOVA followed by Tukey’s multiple-comparison test was carried out, and statistical significance is indicated by *p < 0.05, **p < 0.01, ***p < 0.001.

**Supplementary Information Figure S5.**
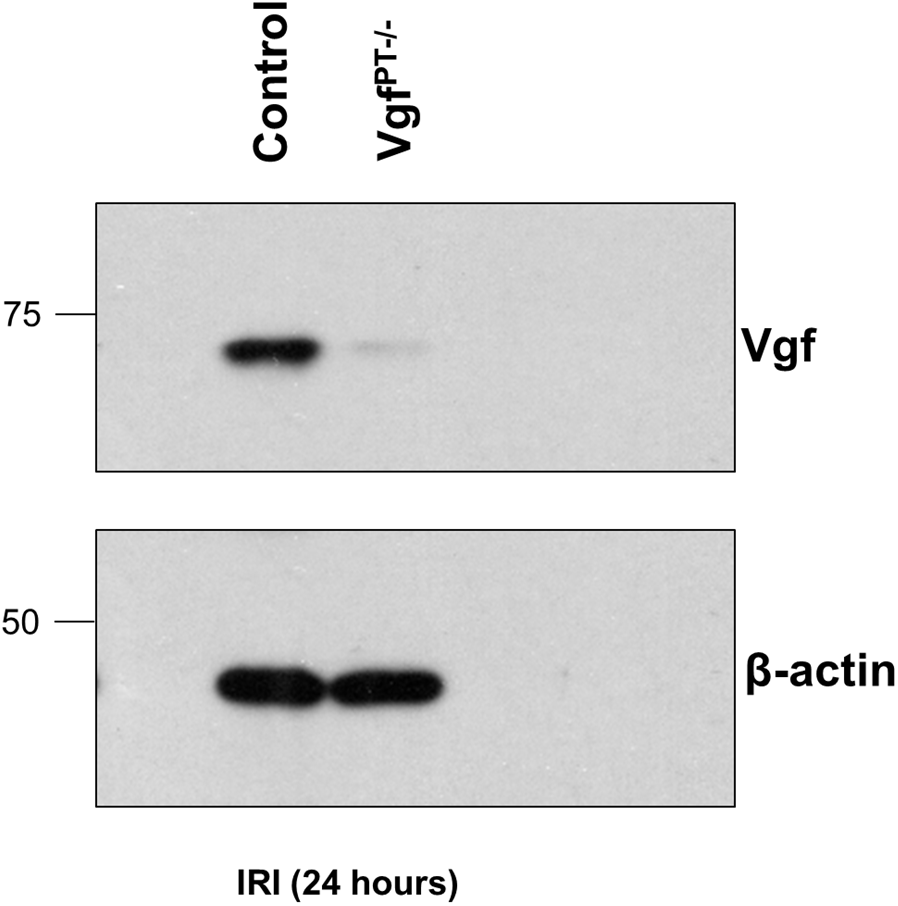
Uncropped Immunoblots for Figure 4J.

